# Assessment of Proton-Coupled Conformational Dynamics of SARS and MERS Coronavirus Papain-like Proteases: Implication for Designing Broad-Spectrum Antiviral Inhibitors

**DOI:** 10.1101/2020.06.30.181305

**Authors:** Jack A. Henderson, Neha Verma, Jana Shen

## Abstract

Broad-spectrum antiviral drugs are urgently needed to stop the COVID-19 pandemic and prevent future ones. The novel severe acute respiratory syndrome coronavirus 2 (SARS-CoV-2) is related to SARS-CoV and Middle East respiratory syndrome coronavirus (MERS-CoV), which have caused the previous outbreaks. The papain-like protease (PLpro) is an attractive drug target due to its essential roles in the viral life cycle. As a cysteine protease, PLpro is rich in cysteines and histidines and their protonation/deprotonation modulates catalysis and conformational plasticity. Here we report the pK_a_ calculations and assessment of the proton-coupled conformational dynamics of SARS-CoV-2 in comparison to SARS-CoV and MERS-CoV PLpros using a newly developed GPU-accelerated implicit-solvent continuous constant pH molecular dynamics method with an asynchronous replica-exchange scheme. The calculated pK_a_’s support the catalytic roles of the Cys-His-Asp triad. We also found that several residues can switch protonation states at physiological pH, among which is C270/271 located on the flexible blocking loop 2 (BL2) of SARS-CoV-2/CoV PLpro. Simulations revealed that the BL2 conformational dynamics is coupled to the titration of C271/270, in agreement with the crystal structures of SARS-CoV-2 PLpro. Simulations also revealed that BL2 in MERS-CoV PLpro is very flexible, sampling both open and closed states despite the lack of an analogous cysteine. Our work provides a starting point for more detailed mechanistic studies to assist structure-based design of broad-spectrum inhibitors against CoV PLpros.

## I. INTRODUCTION

Over the last two decades, three coronaviruses have caused deadly epidemics, threatening the global human population. The severe acute respiratory syndrome coronavirus (SARS-CoV) caused an outbreak in 2003, and a related Middle-East respiratory syndrome coronavirus (MERS-CoV) caused an outbreak in 2012. Today the world is facing the pandemic of the Coronavirus Disease 2019 (COVID-19), caused by a novel coronavirus SARS-CoV-2, which shares about 82% genome sequence identity with the original SARS-CoV^1^. All three viruses are thought to have originated from animal reservoirs and zoonotic transmission into the human population has led to the outbreaks^2^. Currently, no effective treatment exists for any of the three coronavirus diseases; thus, there is an urgent need to understand the potential therapeutic targets and develop inhibition strategies.

Following the release of the coronavirus genome from the acidic endosome, the replicase polyproteins are translated and subsequently self-cleaved by two cysteine proteases to produce the functional non-structural proteins (Nsps) that are required for viral replication. The papain-like protease (PLpro) located in Nsp3 produces Nsp1, Nsp2, and Nsp3, while the 3C-like or main protease located in Nsp5 cleaves 11 sites downstream of Nsp4.^2–6^ In addition to the proteolytic function, CoV PLpro counteracts the host cell innate immune response by deactivating signaling cascades that lead to the impairment of production of pro-inflammatory cytokines and interferons.^7,8^ The former is accomplished through a deubiquitinating (DUB) activity which leads to the removal of ubiquitin from signaling proteins,^9^ and latter through the deIS-Gylating activity which leads to the removal of ISG15 from IRF3.^10^ Thus, PLpro is a critical player in the viral life cycle and as such an attractive drug target for stopping COVID-19 and other coronavirus outbreaks.

Most recently, the first (and only) two X-ray structures of SARS-CoV-2 PLpro were determined (PDB 6W9C and 6WRH, Osipiuk, Joachimiak et al., to be published). The PLpro monomer (about 300 residues), which is the predominant form in solution^11^, is comprised of an independent N-terminal ubiquitin-like (Ubl) domain (first 62 residues) and a C-terminal catalytic domain (Fig. 1a and b). The latter folds in a canonical thumb-palm-fingers like structure, with the Ubl domain anchored to the thumb. The interface between the thumb (residue 107-113, 162-168) and palm (residue 269-279) forms the substrate binding site leading to the catalytic triad of the active site comprised of Cys111, His272, and Asp286 (Fig. 1b). The substrate binding site is solvent exposed and flanked by a flexible *β*-hairpin loop called the blocking loop 2 or BL2 (G266-G271). The fingers subdomain contains a zinc finger coordinated by four cysteines, which upholds the structural integrity and is essential for the PL-pro activity^12^. The structure of SARS-CoV-2 PLpro is nearly identical to that of SARS-CoV PLpro^13^, as expected from the highly similar sequences (96% similarity and 83% identity, Fig. 1c). In contrast, although the structure of MERS-CoV PLpro overlays well with the SARS-CoV PLpro structures, small differences are visible (Fig. 1a), as expected from the larger sequence differences (66% similarity and 30% identity with SARS-CoV PLpro, Fig. 1c).

**FIG. 1.**
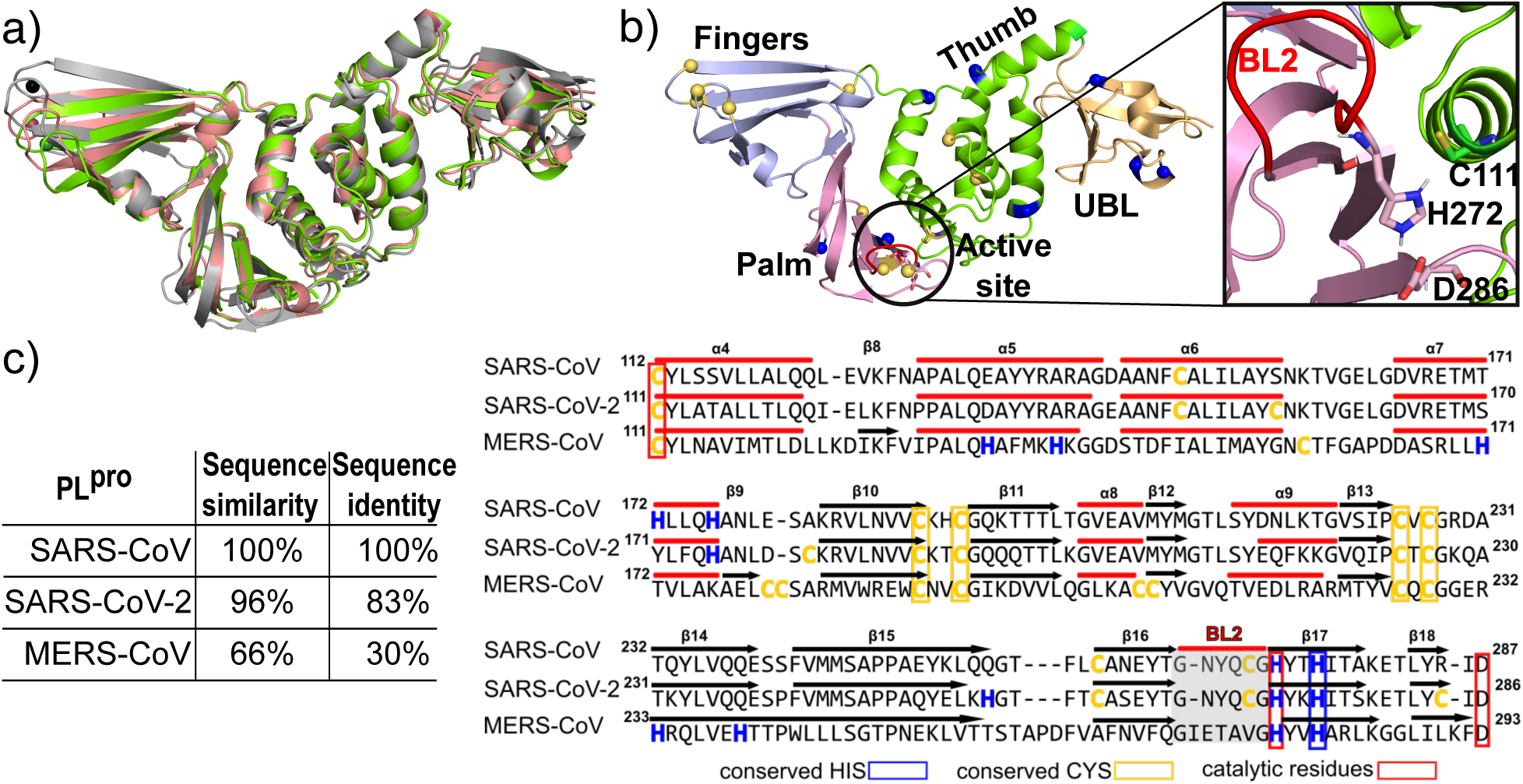
Structure and sequence of SARS-CoV-2 PLpro in comparison to SARS-CoV and MERS-CoV PLpros. a) X-ray crystal structure of SARS-CoV PLpro (in gray, PDB 2FE8^13^), MERS-Cov PLpro (in salmon, PDB 4RNA^24^) overlaid on SARS-CoV-2 PLpro (in green; PDB 6W9C, Osipiuk, Joachimiak et al.,to be published). b) The thumb (residues 63-182), palm (residues 241-314) and fingers (residues 183-240) subdomains of SARS-CoV-2 PLpro are shown in different colors. The Cys and His residues are represented by yellow and blue spheres, respectively. The active site is also shown in a zoomed-in view, with the catalytic His, Cys, and Asp sidechains represented by the stick model and the BL2 loop (G266–G271) colored red. c) Sequence similarity and identity between SARS-CoV, SARS-CoV-2, and MERS-CoV PLpros. The sequence alignment of a part of the thumb and palm subdomains is shown.

All three CoV PLpros are rich in Cys and His residues. SARS-CoV/CoV-2 PLpro contains 8/11 Cys and 11/9 His, while MERS-CoV PLpro contains 14 Cys and 10 His residues (Fig. 1c). Among them, 5 Cys and 2 His residues are conserved among all three PLpros, including the catalytic Cys and His. Cys and His have model pK_a_’s of 8.6 and 6.5, respectively; thus, in model compounds or peptides at physiological pH 7.4, they are both predominantly neutral, i.e., Cys is protonated and His is singly protonated. However, in the protein environment, a small pK_a_ downshift for Cys or up-shift for His may occur, leading to a significant population of or a complete switch to the alternative protonation state, i.e., negatively charged, deprotonated Cys and positively charged, doubly protonated His.

Protonation state switch is an important energy transduction mechanism to enable functionally required conformational changes in biology. For example, the coronavirus spike protein makes use of protonation state switches to induce large conformational changes required for membrane fusion^14,15^. Our previous work demonstrated that proton-coupled conformational dynamics can be studied by continuous constant pH molecular dynamics (CpHMD) simulations^16^ to obtain deeper understanding of the structure-function relationships^17^ and inhibition mechanisms^18–20^ of aspartyl proteases.

Towards understanding the (possibly) proton-coupled structure-function relationship and assisting broad-spectrum inhibitor design, here we report the pK_a_ calculations and preliminary assessment of proton-coupled conformational dynamics of SARS-CoV-2 PLpro in comparison to SARS-CoV and MERS-CoV PLpros. The work employed the recent GPU implementation^21^ of the GB-Neck2 implicit-solvent based CpHMD method^22^ as well as the conventional fixed-charged MD in Amber18^23^. The simulations allowed us to determine the protonation states of all titratable sites, including the catalytic Cys-His-Asp triad, offering a timely knowledge to facilitate MD studies of PLpros in the community. Importantly, we tested a hypothesis regarding the proton-coupled conformational plasticity of the BL2 loop, which modulates substrate and inhibitor binding. The contrasting features among the three CoV PLpros have implications for designing broad-spectrum antiviral inhibitors.

## II. METHODS AND PROTOCOLS

### a. System Preparation

The coordinates were retrieved from the protein data bank (PDB): SARS-CoV PLpro (PDB 2FE8^13^), SARS-CoV-2 PLpro (PDB 6W9C, Osipiuk, Joachimiak et al., to be published), and MERS-CoV PLpro (PDB 4RNA^24^). If multiple chains were available in the x-ray crystal structure, only the first chain was used. Any small molecules or solvent were removed. For each structure, the acetylated N-terminus and amidated C-terminus along with all missing hydrogens were added using the CHARMM program (C36b2)^25^. In the crystal structure of SARS-CoV-2 PL-pro (PDB 6W9C), a disulfide bond is present in the fingers subdomain in place of a zinc ion. Considering that zinc ion cannot be represented in implicit-solvent simulations, we removed the zinc ion and added an analogous disulfide linkage between the closest non-adjacent cysteine pairs in all other structures to maintain the integrity of the fingers subdomain. The disulfide linkage should not affect the pK_a_’s of discussed residues, as they are located far away in other subdomains (Fig. 1a). Following the addition of hydrogens and disulfide bridge, the structure was subject to a 20-step energy minimization with the heavy atoms restrained, and a 20-step energy minimization with all atoms restrained but the disulfide bonded cysteine pairs. The minimization used 10 steps of steepest decent and 10 steps Newton-Raphson methods. From there the force field parameters and coordinate files were constructed from the CHARMM output with the LEAP utility in AMBER.^23^ The ff14sb force field^26^ was used to represent the protein, and the GB-Neck2 (igb=8) implicit-solvent model^27^ was used to represent solvent. The mbondi3 intrinsic Born radii were modified for improving titration simulations of His^28^ and Cys^29^ sidechains. The structure was then energy minimized and equilibrated in the GB-Neck2 implicit solvent^27^, following our previous protocol^21^. The energy minimization was performed using the steepest decent algorithm for 5000 steps and the conjugate-gradient algorithm for 1000 steps. The equilibration was performed at pH 7 in four stages, each having 2000 MD steps with gradually decreased restraining force constants of 5, 2, 1, and 0 kcal/mol/^2^. The final structure was used for CpHMD titration simulations.

### b. pH replica-exchange CpHMD simulations

The titration simulations were performed using the GPU-accelerated GBNeck2-CpHMD method^28,30^ in the pmemd engine of AMBER18.^23^ An asynchronous version (Liu, Harris, Shen, unpublished) of the pH replica-exchange protocol^16^ was implemented to allow replica exchange to be performed on a single GPU card. Our previous work showed that pH replica-exchange enhances both protonation and conformational state sampling, allowing pK_a_’s to rapidly converge.^16,30,31^ In the protocol, 9 pH replicas were placed at pH values ranging from pH 4.5 to 8.5 at an interval of 0.5 pH units. An exchange of two adjacent pH conditions was attempted every 1000 MD steps (or 2 ps). Each replica was run for 55 ns, resulting in an aggregate time of 495 ns for each. The *λ* values were recorded after each exchange attempt. All sidechains of Asp, Glu, His, Cys, and Lys were allowed to titrate, with their titration model parameters taken from our previous work.^28–30^ An ionic strength of 0.15 M was used to represent the physiological salt condition. Simulations were run at a temperature of 300 K and an effectively infinite cutoff (999 Å) for nonbonded interactions. SHAKE was used to constrain bonds involving hydrogens to allow for a 2-fs time step.

### c. Conventional all-atom fixed-charge MD simulations

To support the findings from the GB-CpHMD simulations, two all-atom MD simulations were carried out for SARS-CoV-2 PLpro (PDB 6W9C), using the predicted protonation states for Asp, Glu, His, Cys, and Lys at pH 8.5. Two additional runs were carried out using the protonated form of Cys270. All simulations were performed with Amber18^23^. The protein and water were represented by the ff14SB^26^ and TIP3P^32^ force fields. The initial structure was placed in a truncated octahedron box of water molecules. Long-range electrostatic interactions were handled by using the Particle Mesh Ewald method^33^. A non-bonded cut-off of 8 Å was used with a time step of 2 fs. The starting structure underwent energy minimization by applying 5000 steps of steepest descent, followed by 5000 steps of conjugate gradient minimization with a force constant of 25 kcal/mol/Å^2^ applied to the solute heavy atoms. The force constant was reduced to 5 kcal/mol/Å^2^ and the system was heated from 100 K to 300 K for 50 ps. Following heating, solvent was equilibrated in the NPT ensemble for 250 ps using the isotropic Berendsen barostat^34^ and with the same force constant. Subsequently, the restraints were removed and the system was further relaxed for 100 ps in the NPT ensemble. Finally, two production runs of 1 *µ*s each were performed for each system starting from a different random initial velocity seed. All analysis was performed with the Amber module CPPTRAJ^35^. The first 300 ns from each trajectory was discarded.

## III. RESULTS AND DISCUSSION

We performed pH replica-exchange CpHMD simulations to estimate the pK_a_ values of Asp/Glu/His/Cys/Lys sidechains and assess possible proton-coupled dynamics in SARS-CoV, SARS-CoV-2, and MERS-CoV PLpros. The titration simulations were conducted in the pH range 4.5–8.5 and lasted 55 ns per replica (aggregate simulation time of 495 ns for each protein). The protonation states were well converged. Consistent with our previous work^31^, we found that the protonation states of His residues converge rapidly within 10 ns per replica, and those of Cys converge more slowly due to the formation of hydrogen bonds that are not present in the crystal structure (see later discussion). The time series analysis of protonation state sampling and replica walks along the pH ladder are given in see supplementary material. For the ease of discussion, we refer to the “standard” protonation states as the default settings in the MD programs, i.e., deprotonated Asp/Glu (negatively charged), deprotonated His (neutral), with one proton on either *δ* or *ε* nitrogen, protonated Cys (neutral), and protonated Lys (positively charged). Our simulations showed that several Cys, His residues and one Asp are in the “non-standard” protonation state or can switch to this state at physiological pH in all three PLpros (Table I). A complete list of the calculated pK_a_’s are given in supplementary material.

**TABLE I.**
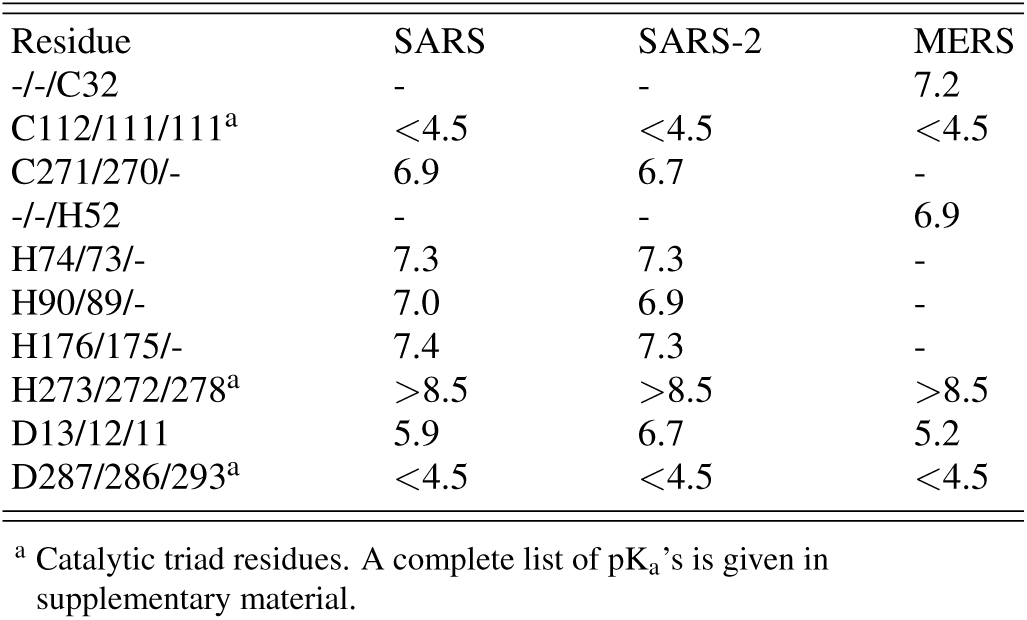
Calculated pK_a_’s of the catalytic residues and those that may switch protonation states at physiological pH in SARS-CoV, SARS-CoV-2, and MERS-CoV PLpros

### a. Protonation states and hydrogen bond network of the catalytic triad

We first consider the catalytic triad in SARS-CoV-2 PLpro. The catalytic Cys111 is located in the thumb, His272 is located in the foothill of the palm adjacent the flexible loop BL2, and Asp286 is located at the end of *β* 18 (Fig. 1b). Currently, no measured pK_a_ data are available. Biochemical experiments of SARS-CoV PLpro suggested that the Cys serves as a nucleophile, while the His functions as a general acid with the assistance of a negatively charged Asp^12^; however, it is unclear whether the reactive nucleophile is the thiolate of the Cys … His ion pair or the neutral thiole which becomes deprotonated upon binding of the substrate^4^. The calculated pK_a_’s of Cys111 and Asp286 are *<*4.5, whereas the pK_a_ of His272 is *>*8.5, indicating that the catalytic triad residues are all in the charged state at physiological pH. Thus, our data supports the mechanism in which the reactive nucleophile is the thiolate ion, rather than the neutral thiol that needs to be first activated by the substrate.^4^

The CpHMD simulations showed that the Cys-His-Asp triad maintains a catalytic geometry through several hydrogen bonds, which support their protonation states. The catalytic His272 forms a hydrogen bond simultaneously with Cys111 and Asp286 in the entire pH range 4.5–8.5, stabilizing His272 in the doubly protonated state and Asp286 and Cys111 in the deprotonated states (Fig. 2a and d). The doubly protonated form allows His272 to perform its role as a general acid.^4^ The catalytic Cys111 forms a hydrogen bond not only with His272 but also with the indole nitrogen of Trp106, which provides further stabilization for the thiolate form and explains the significant downshifted pK_a_ relative to the solution value of 8.5 (Table I). The Cys111 … Trp106 hydrogen bond is important, as it maintains the position of Trp106; the analogous residue in SARS-CoV PLpro has been hypothesized as the oxyanion hole residue that donates a hydrogen bond to stabilize the negatively charged tetrahedral intermediate developed in the course of peptide hydrolysis.^13^ The three hydrogen bond interactions in SARS-CoV-2 PLpro are consistent with those in SARS-Cov PLpro (Fig. 2b).

**FIG. 2.**
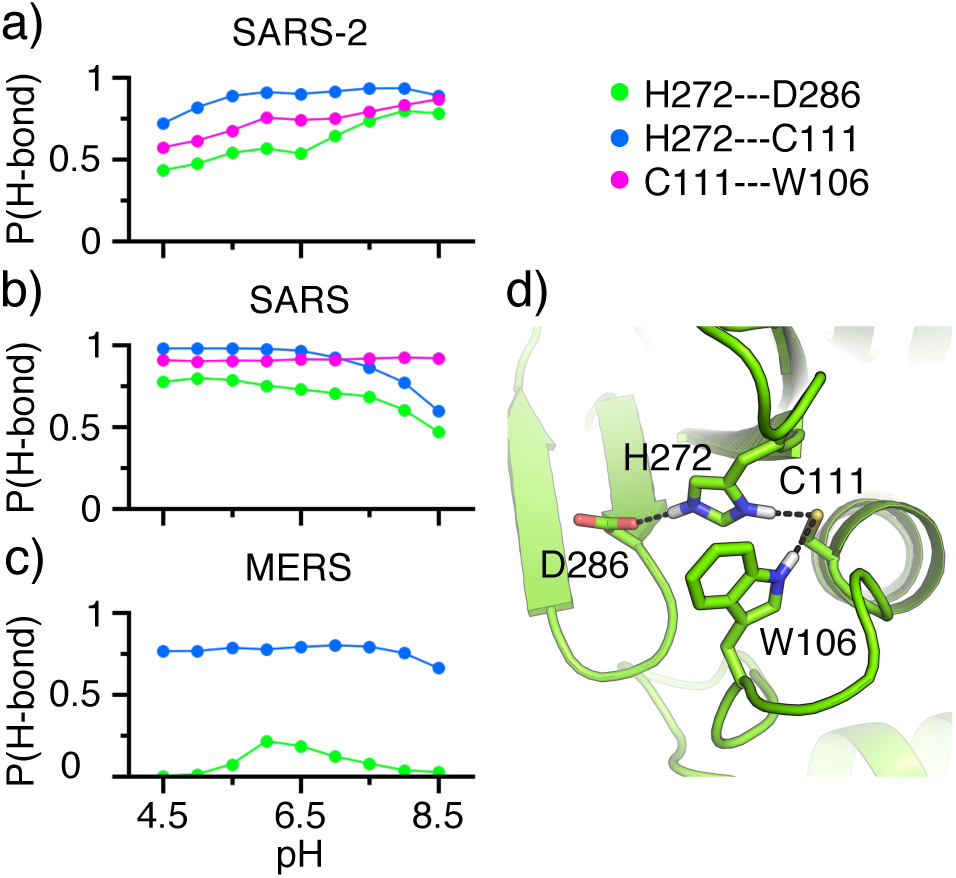
Hydrogen bond formation of the catalytic triad in the PL-pros. a-c) Occupancy of the hydrogen bond between the doubly protonated catalytic His and the deprotonated catalytic Asp (green) or Cys (blue), as well as between Trp106 and the deprotonated catalytic Cys (magenta) as a function of pH in SARS-CoV-2 (top), SARS-CoV (middle), and MERS-CoV (bottom) PLpros. Residue numbering in SARS-CoV-2 PLpro is used. A hydrogen bond was defined using a distance cutoff of 2.4 Å between the hydrogen and oxygen or nitrogen atoms. Data from the last 25 ns/replica were used in the calculations. d) A snapshot showing the hydrogen bonds formed by the catalytic triad in SARS-CoV-2 PLpro.

In MERS-CoV PLpro, Trp106 is replaced with a Leu, which is incapable of forming a hydrogen bond with the catalytic Cys or with the negatively charged intermediate. The has been hypothesized as a cause for the significantly lower catalytic activity of MERS-CoV as compared to SARS-CoV PLpro^36^. In addition to the missing Cys … Trp hydrogen bond, CpHMD simulations of MERS-CoV PLpro showed that the hydrogen bond between the catalytic His and Asp is nearly abolished (Fig. 2c). The loss of hydrogen bond interactions involving the catalytic His and Cys appears to provide less stabilization to the respective charged states in MERS-CoV PLpro, as suggested by the partial titration at the highest and lowest pH condition, respectively (see supplementary material). To test whether the loss of hydrogen bond network affects the flexibility of the regions near the catalytic triad, we calculated the root-mean-square fluctuations (RMSFs) of the C*α* atoms of the catalytic triad and nearby 5 residues (Fig. 3). Interestingly, the RMSFs of the loop residues that are sequence neighbors of the catalytic triad in MERS-CoV PLpro are increased as compared to those in SARS-CoV-2 and SARS-CoV PLpros, which are similar except for the flexible BL2 loop (267–271) region. The loop (106–116) adjacent to *α*4 which harbors the catalytic Cys, the BL2 loop next to the catalytic His, and the *β*-hairpin loop next to the catalytic Asp all display enhanced mobility (Fig. 3a and b). The largest increase in RMSF is seen for the BL2 loop, whereby the RMSF in MERS-CoV PLpro is nearly doubled relative to SARS-CoV-2 PLpro, which shows a somewhat higher mobility than SARS-CoV PLpro. The extremely high flexibility of the BL2 loop in MERS- as compared to SARS-CoV PLpro is consistent with the lack of electron density for the region in the first X-ray structure of the apo MERS-CoV PLpro^36^.

**FIG. 3.**
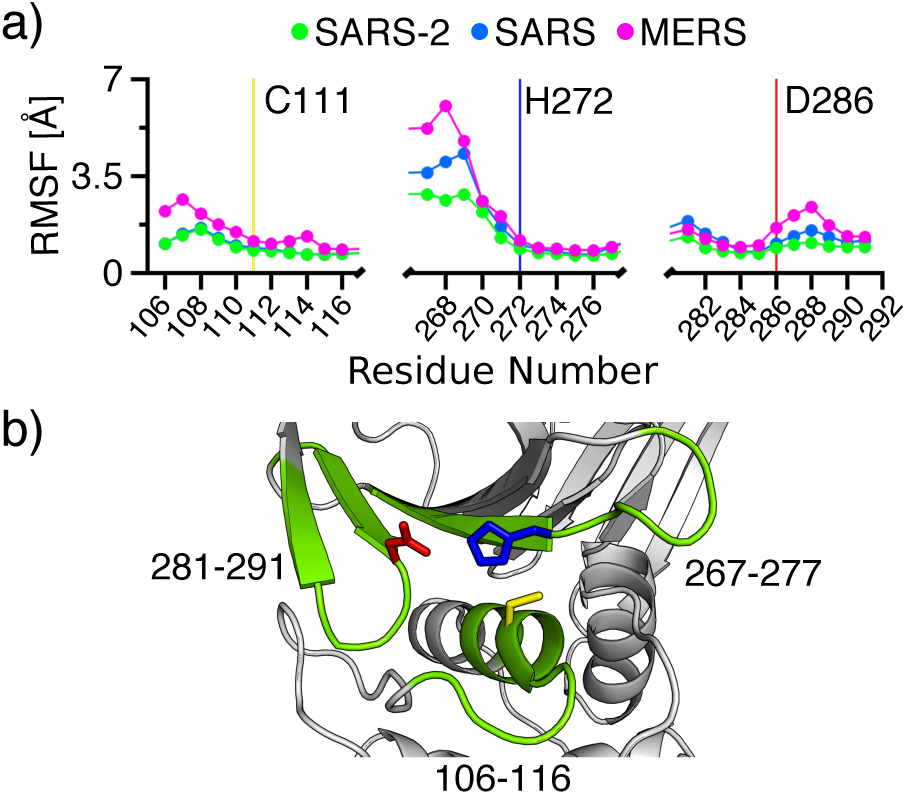
Flexibility of the catalytic triad regions of the PLpros. a) Root-mean-square fluctuations of the C*α* atoms of the catalytic triad and nearby 5 residues in SARS-CoV-2, SARS-CoV, and MERS-CoV PLpros. The sequences of SARS-CoV-2/CoV and MERS-CoV PL-pros are aligned, and the residue numbering of SARS-CoV-2 PLpro is used. b) A snapshot of the catalytic triad regions in SARS-CoV-2 PLpro. Cys111 (yellow), His272 (blue), and Asp286 (red) sidechains are explicitly shown, and the nearby residues are colored green.

### b. Proton-coupled conformational dynamics of the BL2 loop in SARS-CoV/CoV-2 PLpro

The BL2 loop is perhaps the most prominent feature of the substrate binding site in SARS-CoV PLpro, as its movement modulates the substrate and inhibitor binding^4^. Crystal structures show that BL2 is open in the unbound SARS-CoV PLpro and it closes by about 1.5–2 Å in the bound form, which allows hydrogen bonds to form between Tyr269/Gln270 and the inhibitor^4^. Upon inspection of the X-ray structures of SARS-CoV-2 PLpro, We noticed that the BL2 loop is open in the pH 7.5 structure (PDB 6W9C) and a zinc ion is found within the binding distance of Cys270; however, in the structure determined at pH 4.5 (PDB 6WRH), the BL2 loop closes in by 1.9 Å (C*α* distance between Tyr268 and Asp164) and the zinc ion is absent. Thus, we hypothesized that the BL2 loop dynamics is coupled to the titration of C270.

CpHMD titrations gave the pK_a_’s of 6.7 and 6.9 for Cys270 in SARS-CoV-2 PLpro and the equivalent Cys271 in SARS-CoV PLpro, respectively (Table I). Thus, Cys270/C271 samples both protonated and deprotonated states at physiological pH. The pK_a_ downshifts relative to the model Cys pK_a_ of 8.5 is due to the formation of local hydrogen bonds which favors the thiolate state. In the crystal structure of SARS-CoV-2 PL-pro, Cys270 does not interact with Thr265. The titration simulations showed that the distance between Cys270 and Thr265 varies widely between 5 and 15 Å, when Cys270 is protonated (*λ* value close to 0); however, when Cys270 is deprotonated (*λ* value close to 1), the distance is locked to 1.5–2.5 Å, indicating the formation of a hydrogen bond (Fig. 4a). In addition to the hydroxyl group of Thr265, the deprotonated Cys270 can also accept a hydrogen bond from the backbone amide groups of His272 and Gly271, stabilizing the charged state (Fig. 4b). The same hydrogen bonds were also formed in the simulations of SARS-CoV PLpro, which explains the similarly downshifted pK_a_ of the equivalent Cys271.

**FIG. 4.**
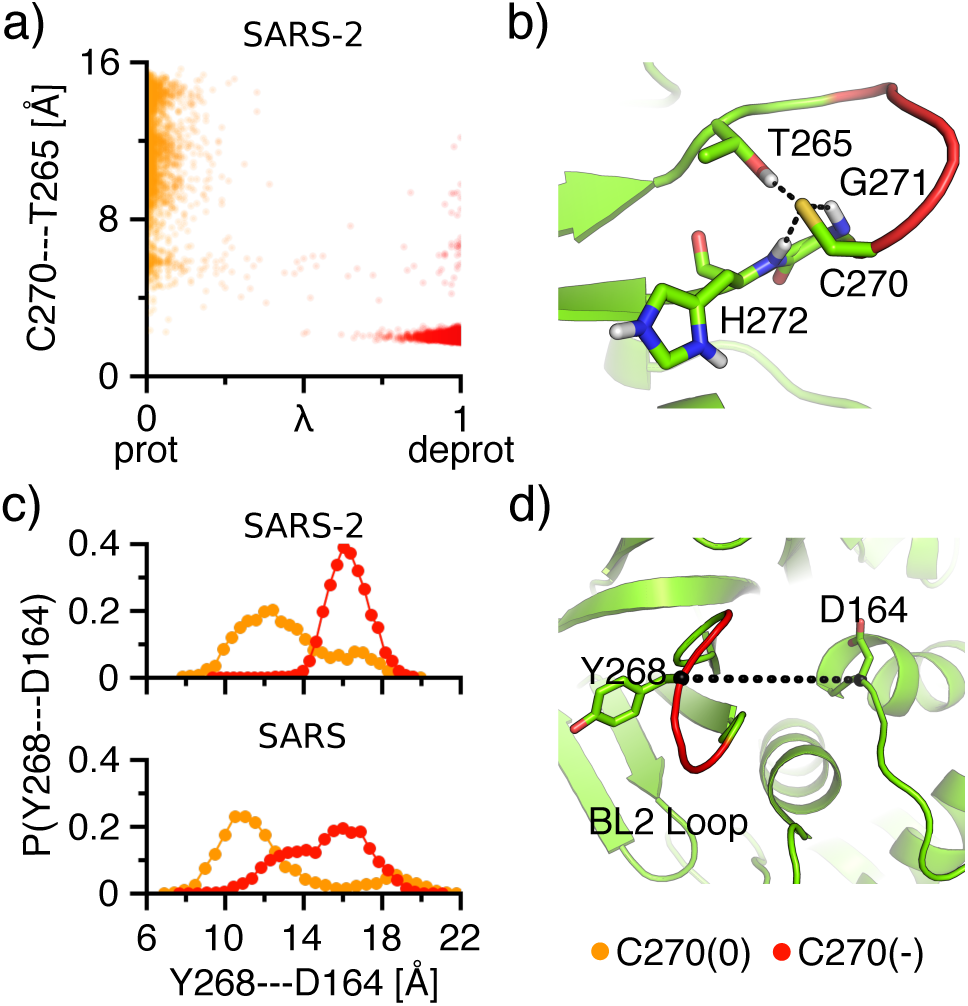
Titration of Cys270 is coupled to the conformational dynamics of the BL2 loop in SARS-CoV/CoV-2 PLpro. a) Correlation between the protonation state of Cys270 and the distance between the sulfur of Cys270 and the hydroxyl hydrogen of Thr265 in the SARS-CoV-2 PLpro simulation at pH 7. *λ <*0.2 (orange) and *λ >*0.8 (red) are used to define protonated and deprotonated states, respectively. b) A snapshot showing the BL2 loop (red) and the hydrogen bonds formed around a deprotonated Cys270 in SARS-CoV-2 PLpro. c) Probability distributions of the C*α* distance between Tyr268 and Asp164, when Cys270 is protonated (orange) or deprotonated (red) from the simulations of SARS-CoV-2 (top) and SARS-CoV (bottom) PLpro at pH 7. Data from pH 7.5 and pH 8 simulations are similar and not shown here. d) A snapshot showing the BL2 loop environment with Y268 and D164 labeled.

To test the hypothesis that protonation/deprotonation of Cys270 modulates the BL2 dynamics, we examined the C*α* distance between Tyr268 on the BL2 loop and the conserved Asp164 next to *α*7, which represents the width of the S3 subpocket (Fig. 4c and d). The equivalent Tyr269 in SARS-CoV PLpro is an important residue, as it forms a hydrogen bond with the inhibitors that bind to the S3 pocket^4^. When Cys270 is protonated, the probability distribution of the Tyr268–Asp164 distance covers a broad range of 8–20 Å with a peak at around 12 Å; however, when Cys270 is deprotonated, the distribution samples a narrower range of 14–20 Å with a peak at around 16 Å (Fig. 4c). Thus, the CpHMD data suggests that the deprotonated Cys270 is correlated with the BL2 conformations that are more open and rigid, which might be attributed to the aforementioned hydrogen bond formation between the deprotonated Cys270 and the surrounding residues. Turning to SARS-CoV PLpro, Fig. 4c shows that the BL2 movement is coupled to the protonation/deprotonation of the analogous Cys271. However, in SARS-CoV PLpro, it appears the BL2 with a deprotonated Cys270 can sample a wider range of 10–20 Å as compared to SARS-CoV-2 PLpro, although the peak remains around 16 Å. The wider range of BL2 movement may be attributed to the somewhat weaker hydrogen bonds involving the deprotonated Cys271. The movement of the BL2 loop is very similar between SARS-CoV and SARS-CoV-2 PLpros when Cys270 is protonated.

### c. Comparison of the BL2 conformation across the three PLpros

For broad-spectrum inhibitor design, it is important to understand the difference in the BL2 conformation across the three PLpros. The distributions of the Tyr269/268– Asp165/164 distance for SARS-CoV/CoV-2 and the equivalent Thr274–Asp164 distance for MERS-CoV PLpro at physiological pH (Fig. 5a and Fig. 4d) show that the widest position of the BL2 loop is about the same across the three PLpros; however, the BL2 in SARS-CoV-2 PLpro samples the narrowest range, followed by SARS-CoV PLpro and MERS-CoV PLpro which samples the widest range between 5–22 Å. The enhanced flexibility of the BL2 in MERS-CoV PLpro may be attributed to the lack of a Cys equivalent to Cys271/270 in SARS-CoV/CoV-2 PLpro which can form hydrogen bonds with neighboring residues to restrict the loop motion and perhaps also the loosening of the nearby catalytic His which no longer forms double hydrogen bonds as in SARS-CoV/CoV-2 PLpro.

**FIG. 5.**
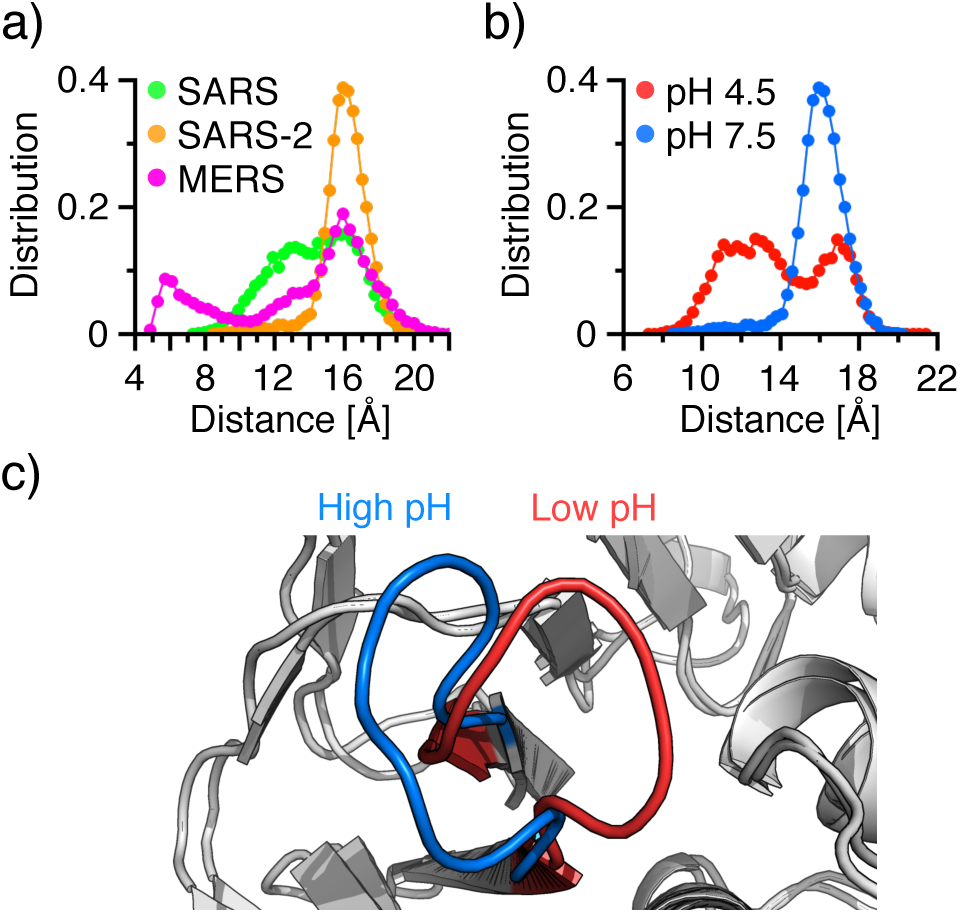
Comparison of the BL2 loop conformation across the three PLpros and between low and high pH. a) Probability distributions of the C*α* distance between Tyr269/268 and Asp165/164 in SARS-CoV/CoV-2, and between Thr274 and Asp164 in MERS-CoV PLpro at pH 7.5. b) Probability distribution of the C*α* distance between Tyr268 and Asp164 in SARS-CoV-2 PLpro at pH 4.5 (red) and 7.5 (blue). c) Overlaid crystal structures showing the closed (red; pH 4.5 structure) and open (blue; pH 7.5 structure) BL2 conformations.

In agreement with the BL2 dynamics described with the protonated and deprotonated states of Cys270 (Fig. 4c) and the X-ray structures determined at pH 7.5 (PDB 6W9C) and pH 4.5 (PDB 6WRH), a significant pH dependence is observed with the BL2 dynamics in the simulations of SARS-CoV-2 PLpro (Fig. 5). In the simulation at pH 7.5, the BL2 loop samples a state that leaves the S3/S4 subpocket more open, similar to the neutral pH crystal structure (Fig. 5c). In the simulation at pH 4.5, the BL2 loop assumes a wide range of dynamics, allowing it to sample a state that leaves the S3/4 subpocket more closed, similar to the low pH crystal structure (Fig. 5c).

### d. The Ubl domain contains an Asp with a highly upshifted pK_a_

As expected, nearly all Asp/Glu residues adopt standard protonation states (i.e. charged) at physiological pH; however, Asp12 in SARS-CoV-2 PLpro has a pK_a_ abnormally upshifted from its model pK_a_ of 4.0 to 6.7 (Table I), making it possible to occasionally sample the protonated state at pH 7.4. Trajectory analysis suggested that this upshift is in part due to the protonated Asp12 acting as a hydrogen bond donor to either (deprotonated) Glu67 or Asn15 (Fig. 6a and c), which stabilizes the protonated state. The two hydrogen bonds are mutually exclusive such that Asp12 is a hydrogen bond donor 82–96% of the time when it is in the protonated state (Fig. 6a). In addition to hydrogen bonding, Asp12 is buried in a hydrophobic pocket with a very low solvent accessible surface area, which increases as Asp12 becomes deprotonated at higher pH (Fig. 6b). In SARS-CoV PLpro, the analogous Asp13 experiences a similar degree of hydrogen bonding and solvent sequestration, resulting in a upshifted pK_a_ of 5.9. In MERS-CoV PLpro, the analogous Asp11 primarily donates a hydrogen bond to Asn15, as the analogous residue to Glu67 is missing (see supplementary material). Compared to Asp12/13 in SARS-CoV-2/CoV PLpro, Asp11 is more solvent exposed in the lower pH range (see supplementary material), which may contribute to a smaller degree of pK_a_ upshift of Asp11 in MERS-CoV PLpro as compared to SARS-CoV/CoV-2 PLpro.

**FIG. 6.**
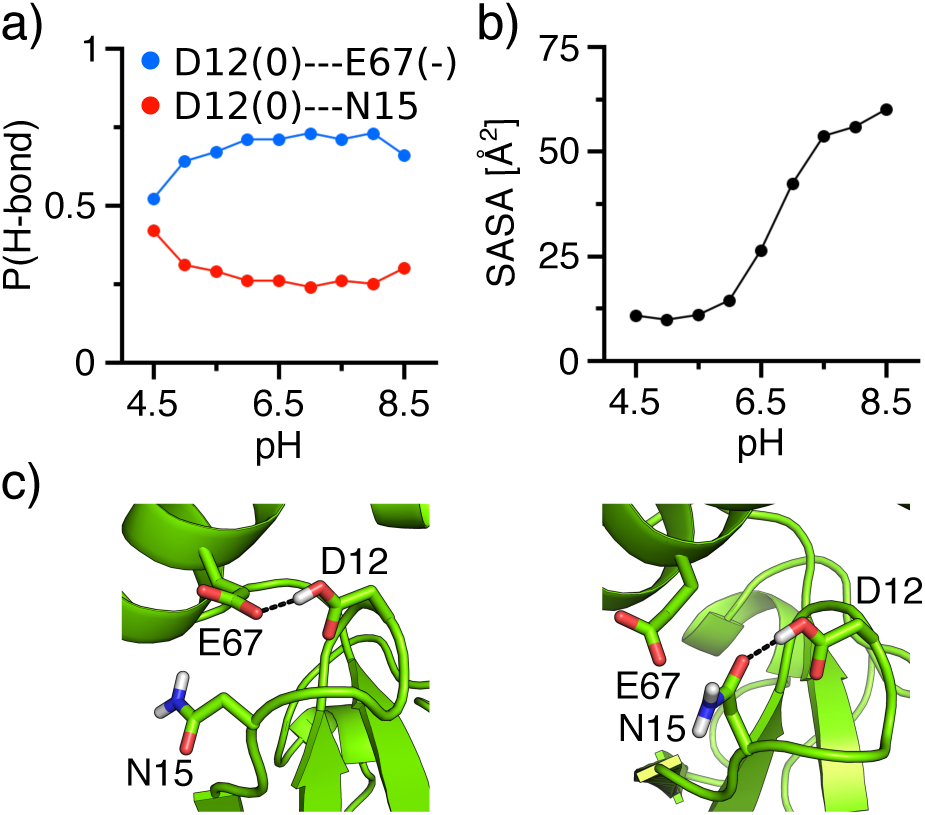
Molecular determinants of the large pK_a_ upshift of Asp12 in SARS-CoV-2 PLpro. a) Occupancy of the hydrogen bond between the protonated Asp12 and the deprotonated Glu67 or Asn15 in SARS-CoV-2 PLpro as a function of pH. b) Solvent accessible surface area of Asp12 (based on the heavy atoms) as a function of pH. c) Snapshots showing the hydrogen bond between Asp12 and Glu67 or Asn15.

### e. Histidines that can switch protonation states at physiological pH

CpHMD titrations revealed that three histidines unique to SARS-CoV/CoV-2, H74/73, H90/89, and H176/175 (Fig. 7), have pK_a_’s around 7 (Table I) and can sample both protonated and deprotonated states at physiological pH. His74/73 located on the C-terminal end of *α*2 in SARS-CoV/CoV-2 has a pK_a_ of 7.3/7.3. Analysis suggested that the pK_a_ upshift relative to the model value of 6.5 is due to the formation of hydrogen bonds with Phe70/69, Asn129/128 or a salt bridge with Glu71/70 (see supplementary material), which stabilizes the charged state. His90/89 located on the C-terminal end of *α*3 in SARS-CoV/CoV-2 has a pK_a_ of 7.0/6.9. Analysis showed that a small pK_a_ upshift relative to the model pK_a_ is due to the stabilization of the charged state by the occasional hydrogen bonding with the backbone carbonyl of Ser86/85 or transient salt-bridge interaction with Asp107/108 located near the oxyanion hole of the CoV PLpro (see supplementary material). His176/175 is located on the C-terminal end of *α*7 and opposite to His74/73. The increased pK_a_ of 7.4/7.3 can also be attributed to local hydrogen bonding, either with His172 in SARS-CoV-1 or Tyr171 in SARS-CoV-2 PLpro (see supplementary material).

**FIG. 7.**
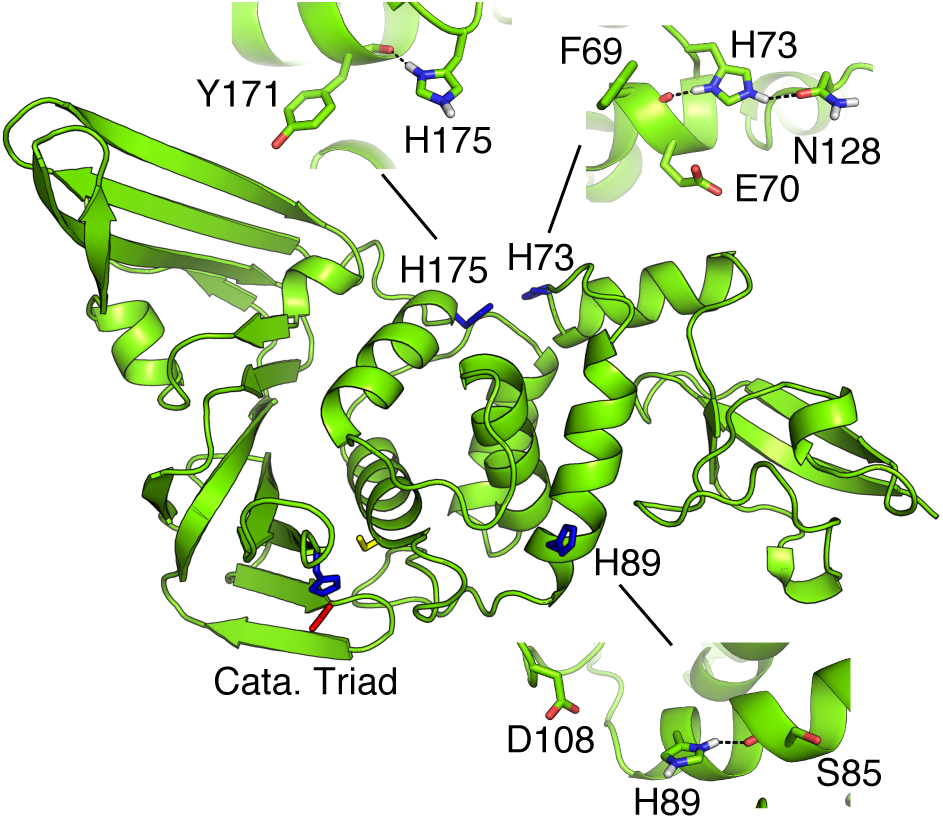
Locations of the three histidines in SARS-CoV/CoV-2 PLpro that can switch protonation states at physiological pH. Residues that provide interactions to stabilize the imidazolium form are shown.

### f. All-atom fixed-charge MD of SARS-CoV-2 PLpro

To provide support for the protonation states determined by GB-CpHMD titrations and test the proton-coupled dynamics of the BL2 loop, we performed conventional all-atom fixed-charge MD simulations of SARS-CoV-2 PLpro with the catalytic sidechains fixed in the charged states and Cys270 fixed in the protonated or deprotonated state. All other residues were fixed in the standard protonation states. Two 1-*µ*s trajectories were obtained with each Cys270 protonation state. Consistent with the GB-CpHMD simulations at physiological pH, the catalytic triad remained very stable, with the hydrogen bond between His272 and Cys111 being the strongest, followed by the His272 … Asp286 and Cys111 … Trp106 hydrogen bonds, as shown in the hydrogen bond occupancy plots (Fig. 8a). Interestingly, while the effect of Cys270 protonation/deprotonation appears negligible for the latter two hydrogen bonds (occupancy change is below 5%), protonation of Cys270 weakens the His272 … Cys111 hydrogen bond (occupancy decreases by over 20%). This decrease is consistent with the GB-CpHMD data, which shows that the catalytic His…Cys hydrogen bond is significantly weakened below pH 6 as Cys270 becomes fully protonated in SARS-CoV-2 PLpro (Fig. 2a).

**FIG. 8.**
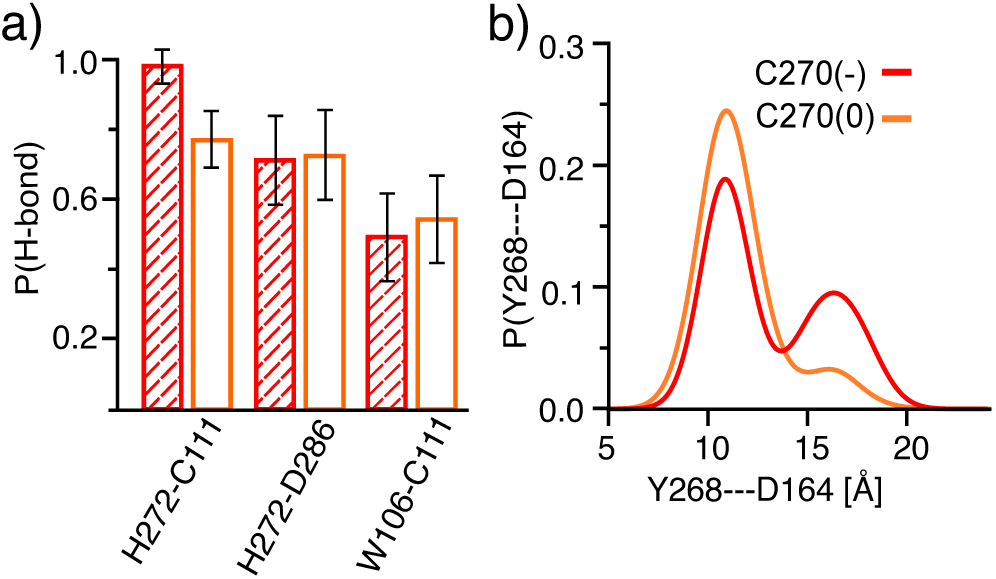
Conventional MD of SARS-CoV-2 PLpro with Cys270 fixed in the protonated or deprotonated state. a) Occupancies of the His272 Asp286, His272 … Cys111, and Cys111 … Trp106 hydrogen bonds from the simulations with Cys270 fixed in the protonated (orange) or deprotonated (red) state. The calculations combined the data from two independent 1-*µ*s trajectories. The first 300 ns data were discarded. b) Probability distribution of the C*α* distance between Tyr268 and Asp164 from the simulations with Cys270 fixed in the protonated (orange) or deprotonated (red) state.

To test the effect of Cys270 titration on the BL2 conformation, we calculated the probability distribution of the C*α* distance between Tyr268 and Asp164. For the trajectories with protonated Cys270, the distribution displays a single peak at about 11 Å; however, for the trajectories with deprotonated Cys270, a second peak appears at about 17 Å (Fig. 8b). This data indicates that deprotonated Cys270 is correlated with the more open BL2 loop conformations, consistent with the findings from the GB-CpHMD simulations (Fig. 4c). However, due to the slow transition between the open and closed BL2 loop conformations in explicit solvent, more trajectories or longer simulations are needed to solidify the conclusion.

## CONCLUDING DISCUSSION

The protonation states and possible proton-coupled conformational dynamics of SARS-CoV-2 PLpro were investigated in comparison to SARS-CoV and MERS-CoV PLpros, using the GPU-accelerated GBNeck2-CpHMD titration simulations with a new asynchronous pH replica-exchange scheme as well as conventional all-atom MD. The simulations showed that the catalytic Cys, His, and Asp are charged in the entire simulation pH range of 4.5–8.5 for all three PLpros, which supports the mechanism in which the reactive nucleophile is the thiolate ion and the catalytic His serves as a general acid stabilized by the catalytic aspartate. The catalytic triad in SARS-CoV-2/CoV PLpro forms a hydrogen bond network among themselves and with a nearby Trp which serves as an oxyanion hole residue to stabilize the tetrahedral intermediate developed in the peptide hydrolysis. In contrast, the hydrogen bond with Trp is missing and the hydrogen bond between the catalytic His and Asp is nearly abolished in MERS-CoV PLpro, consistent with the significantly lower catalytic activity compared to SARS-CoV PLpro^36^. Interestingly, the lack of a hydrogen bond network for the catalytic triad in MERS-CoV PLpro is correlated with increased mobility of nearby loop residues, particular the BL2 loop.

The simulations revealed that several titratable residues have shifted pK_a_ values such that they switch between two protonation states at physiological pH. These include three His and one Cys residues unique to SARS-CoV-2/CoV and one Asp residue common to all three PLpros (Table I). Of particular interest is Cys270/271 on the flexible BL2 loop of SARS-CoV-2/CoV, which has a pK_a_ of 6.7/6.9 and samples both the standard thiol and charged thiolate forms at neutral pH. CpHMD simulations showed that the BL2 loop samples an open or a closed conformational ensemble with deprotonated or protonated Cys270/271, respectively, consistent with two crystal structures of SARS-CoV-2 PLpro determined at neutral and low pH conditions and the conventional all-atom MD trajectories of SARS-CoV-2 PLpro with either deprotonated or protonated Cys270. Thus, the simulation data and experiment together support our hypothesis that the BL2 loop conformation is coupled to the titration of C270/271 in SARS-CoV-2/CoV PLpro.

An induced fit mechanism, by which BL2 closes in to form hydrogen bonds with the inhibitor, has been proposed in designing potent inhibitors targeting the S3/S4 pocket of SARS-CoV PLpro^4^. Our finding suggests that in the absence of a ligand, protonation of C270/271 induces the closure of BL2 in SARS-CoV-2/CoV PLpro, which raises the possibility that a conformational selection mechanism may be operative, in which inhibitor binding shifts the BL2 conformational population to the closed form, perhaps by favoring the protonation of C270/271. While the unliganded crystal structures show that BL2 in MERS-CoV PLpro is more open than in SARS-CoV-2/CoV PLpro, CpHMD simulations suggest that BL2 has an increased flexibility in MERS-CoV PLpro and it can sample closed conformations. This finding challenges a current hypothesis, according to which SARS-CoV PLpro inhibitors do not bind to MERS-CoV PLpro due to the more open BL2 loop as a result of the sequence difference and one extra residue^24,37^. A main caveat of our work is the use of the GB-Neck2 implicit-solvent model^27^. Although it has been demonstrated in the accurate de novo folding simulations of nearly two dozen small proteins with *α* and *β* topologies ^38^, inherent issues such as the lack of solvent granularity may limit the accuracy of detailed conformational representation. Nonetheless, our work provides a starting point for further mechanistic investigations using higher-level approaches such as the all-atom CpHMD^39^ and more extensive conformational sampling to assist the structure-based drug design targeting the coronavirus PLpros.

## SUPPLEMENTARY MATERIAL

See supplemental material for convergence analysis, a list of pK_a_ estimates for all titratable residues, and additional hydrogen bond analysis.

## ACKNOWLEDGMENTS

We acknowledge final support from the National Institute of Health (GM098818).

## DATA AVAILABILITY

The data that support the findings of this study as well as raw trajectory files are available from the corresponding author upon reasonable request.

